# High Temporal Sub-millisecond Time Resolution Stimulation Increases Performances of Retina Prosthetic Vision

**DOI:** 10.1101/442269

**Authors:** S. Kime, C. Simon-Shane, Q. Sabatier, F. Galluppi, R. Benosman

**Affiliations:** Sorhonne Universitas, Institut de la Vision, 17 rue Moreau, F-75012 Paris, France; University of Pittsburgh Medical Center, Eye and Ear Institute, Biomedical Science Tower 33501 Fifth Ave, Pittsburgh, PA 15213, Unites-States; Carnegie Mellon University, Robotics Institute, 5000 Forbes Avenue, PA 15213, Unites-States

## Abstract

Vision loss has an enormous impact on affected individuals, and increasing social and economics costs for the whole of society. Important approaches have been introduced, aimed at restoring visual functionality in the case of vision degradation and loss. Currently the most promising technology is represented by retinal implants, with companies producing devices approved for commercialization. The first clinical studies encouragingly reported perception elicited by both devices. Some patients retrieve autonomy, but only a few could reach satisfactory perception levels, allowing more complex tasks such as reading. Fundamental limitations however still exist in both spatial and temporal resolution, and these limitations call for improvements of the technology and of the stimulation strategies used therein. This study focuses on the importance of temporal resolution for retinal stimulation, and how it can be used to improve restored perception for implanted patients. We show a quantification of the improvement of discrimination performance when stimulating with high frame rates compared to conventional 30-60Hz frame rates. Such effect is evaluated using data collected from implanted patients, and deriving from it a realistic phosphene model. This study allowed to accurately replicate the perceptual effects of the studied technologies, and to evaluate the hypotheses regarding performance improvements using higher temporal resolutions for stimulation strategies in visual restoration devices.

## 1 Introduction

Important research efforts are underway in the field of visual prosthetics to restore visual functionality and thus to increase the autonomy of blind or visually impaired people. Implantable visual prosthetics aim at restoring vision in patients affected by degenerative diseases including retinitis pigmentosa (RP) or aged-related macular degeneration (AMD). Such diseases currently affect around 20 million individuals, including more than 1 million affected with RP [1, 2].

As both diseases affect the visual system at the retinal level, the main strategy to restore vision consists in developing implants made of active or passive electrodes to stimulate the healthy cells in the retina. The prosthesis is called an *epiretinal* implant, when these electrodes are placed on top of the inner surface of the retina. When the implant is placed between the retina and the retinal pigment epithelium (RPE) it is called a *subretinal* prosthesis, in this case the electrodes are small photodiodes which convert light into current that stimulates the healthy layers of the retina [3].

The last decade has seen a significant maturation of the field of prosthetic vision with the development and validation of several devices. As an example, Second Sight commercialized Argus II is an epiretinal retinal implant composed of 60 electrodes of 200 *μm* stimulating a visual field of 10 × 20 deg [4, 5]. Intelligent Medical Implants (IMI) also developed an implant (IRIS) using a similar stimulation technique with 49 electrodes and a field of view up to 40 deg [6, 7]. This implant was tested on 20 patients and the technology has been acquired by Pixium Vision. The second generation of the implant, which comprises 150 electrodes, is undergoing clinical trials [8].^1^

Other companies proposed different design and stimulation techniques. For example Alpha-IMS uses a subretinal implant with 1500 independent micro-photodiodes, capable of stimulating an area of 11 × 11 deg [9, 10]. Likewise, Pixium Vision’s third generation implant (PRIMA) will propose a matrix of micro-photodiodes based on [11]. Bionic Vision has introduced a suprachoroidal retinal prosthesis, and performed clinical trials of a prototype version with 12 electrodes [12, 13]; they are currently developing a prototype with 44 electrodes, a Wide-View device with 98 electrodes, and a High-Acuity device with 256 electrodes.^2^

While such prostheses are capable of restoring visual functionalities up to a degree of success [4, 12, 10], some limitations persist. One of the most concerning limitation is the number of implantable electrodes, which is bounded by the size of the electrodes involved. One area of improvement is in increasing the spatial resolution, so as to improve the quality of visual perception (phosphenes). This involves reducing the size of the electrodes [3], or modifying their shape (3D electrodes) [14]. While introducing some improvements in stimulation, these methods are still confronted with technical and physical limits [15]. Due to the technological issues involved in producing smaller electrodes, other approaches are aiming to improve the restored visual perception. Given the limitation in spatial resolution, in this work we address the possibility of leveraging the timing properties of the visual system to deliver better visual stimulation.

The human visual system has high temporal dynamics: retinas can process information with a resolution of the order of the millisecond. Such high timing precision can be found for example at the ganglion cell level, as shown by Gollisch *et al.* [16, 17] or even in cells located in the visual cortical area MT [18]. At a higher level, the effect of high refresh rates on the perception of moving stimuli has also been demonstrated [19].

In this paper we present a study of the impact of high temporal resolution on spatial acuity with visual prosthetics, aimed at improving restored vision in implanted patients. The goal of our study is to devise novel stimulation strategies and methods, which can guide the design of the next generations of portable visual interfaces to be paired with retinal implants.

Based on data describing typical phosphenes and their probabilities, we devise a phosphene model, described in section 2.2. Phosphenes are represented by thresholded 2D Gaussians to obtain a binary model.

To cope with the reduced spatial resolution and the fact that the camera providing visual input is fixed, implanted patients often use head movements to scan the scene and acquire more information. This needs to be taken into account when devising a phosphene model. Section 3 reports the results and velocity of head movements of an implanted patient performing a visual discrimination task.

Section 4 introduces a stimulation platform capable of delivering visual information in form of precise, high-temporal resolution light pulses. This platform is designed to elicit visual sensations in subjects with normal vision that mimic those described by implanted patients.

We use this platform and the phosphene model to compare different stimulation strategies controlling temporal resolution on ten normally sighted subjects. Subjects performed a series of tasks that are commonly used in psychophysical studies to evaluate the visual acuity and the success rate of natural tasks such as digit recognition We show that temporal resolution can indeed be used to improve retinal stimulation in common visual tasks, as presented in Section 5. The results of our experiments demonstrate the importance of precise timing in delivering a better perception to implanted patients.

## 2 Phosphene model

In prosthetic vision research, the percepts of light induced in implanted patients through means of electrical stimulation of the visual pathway using the devices described in Section 1 are called *phosphenes.* For a review of different phosphene descriptions and models see [20]. To correctly evaluate the efficiency of a stimulation strategy for an implanted patient, we use an accurate phosphene model, based on data from observed phosphenes. This is a challenge, since very little experimental data characterizing phoshenes is available: there are very few implanted patients, and the data collected during clinical trials is proprietary. This section describes typical phosphene characteristics as provided by Pixium Vision based on their clinical trials, and the derived phosphene model. In contrast with most phosphene models that rely on numerous assumptions, ours based on quantitative phosphene characteristics.

### 2.1 Phosphene description

The phosphenes are described qualitatively by their shape, orientation, size, color and brightness. Each of these characteristics is described quantitatively or, when this is not possible, qualitatively in different categories, based on Pixium’s experience: *shape* is either round, oblong or lank; *orientation,* for non round phosphenes, is given as a multiple of 15°; *size* of the phosphene as if it was perceived at arm’s length; *color* chosen from red, blue, yellow, white, green and black; *brightness* is qualitatively described as darker than background, light bug, candle, light bulb or sun.

Typical phosphenes are shown in Figure 1. The phosphenes are represented by a 2D Gaussian which is scaled in intensity depending on the brightness. To allow the representation of the very rare phosphenes that are darker than the background, the background is set to a non-null grayscale (typically 0.1 in our images).

**Figure 1:**
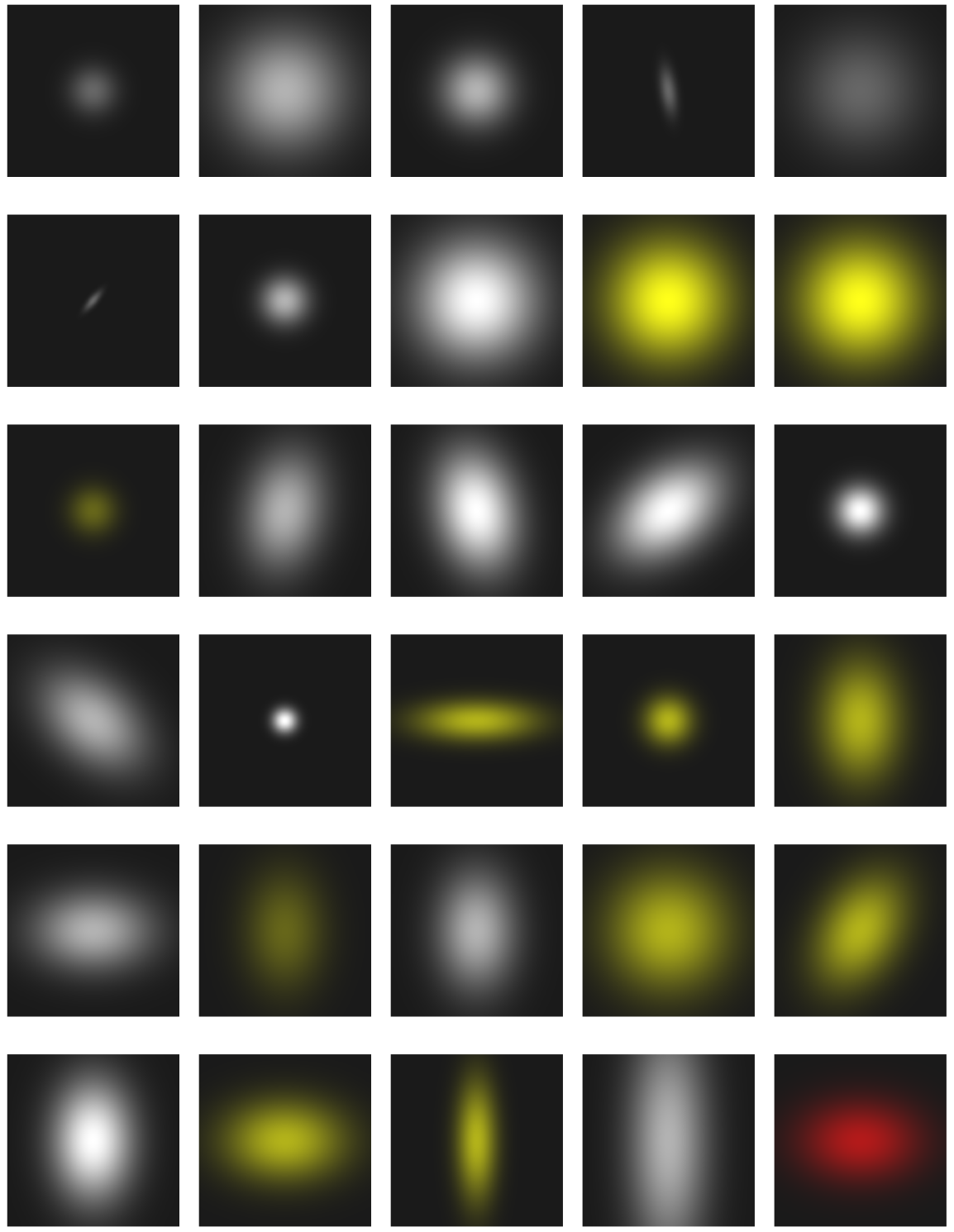
Typical phosphenes. The side of each figure represents 7 cm at arms’ length.

The size of the phosphenes is controlled by varying the standard deviation *σ*_*x*_. The shape of the phosphenes is controlled by varying the relative values of the standard deviation in *x* and *y* (*σ*_*x*_ and *σ*_*y*_). A round phosphene is modeled with *σ*_*x*_ = *σ*_*y*_; an oblong phosphene with 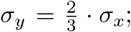 and a lank phosphene with 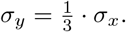.

### 2.2 Phosphene model

We simplify the model to only take into account the size, shape and orientation. This model will be used to project phosphenes directly into subjects eyes, as explained in detail in section 4.2. Since the platform can only project binary data, we model binary phosphenes, without taking into account neither color nor brightness. However, our data shows little variability in the color of the phosphenes perceived: mostly white and yellow phosphenes.

We threshold the output of the Gaussian model used for the phosphene representation. The standard deviation *σ*_*x*_ is chosen so that thresholding at value *T* gives a phosphene of the desired size *S*:

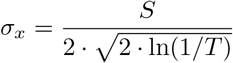

The bandwidth of our setup induces a compromise between the resolution of the largest phosphenes and the size of the phosphene map. We fix the map size to 20 × 25 phosphenes, thus 500 different perceptual sites, and the resolution of the largest phosphenes to 7 pixels. We model the three shapes with their respective probability, accordingly to Pixium’s information. As the data we have shows no preferred orientation, we assume an equal probability of horizontal and vertical non-round phosphenes. Likewise, a majority of phosphenes are described as between 2 cm and 5 cm at arm’s length. We thus model two sizes “Large” and “Small” which respectively correspond to a size of 5 cm represented by 7 pixels and 2 cm represented by 3 pixels. The phosphenes overlap with 4/7 of the width of the largest phosphenes. We assume a perfect retinotopic arrangement. Though this hypothesis is quite strong, it is corroborated by first descriptions of phosphenes [21, 22].

The ten binary 7 × 7 phosphenes and their corresponding probability of occurrence are shown in figure 2. Phosphene maps that respect these statistics are shown in figure 3. The maps are shown with no overlap so that the individual phosphenes can be distinguished, however we use them with a 4/7 overlap.

**Figure 2:**
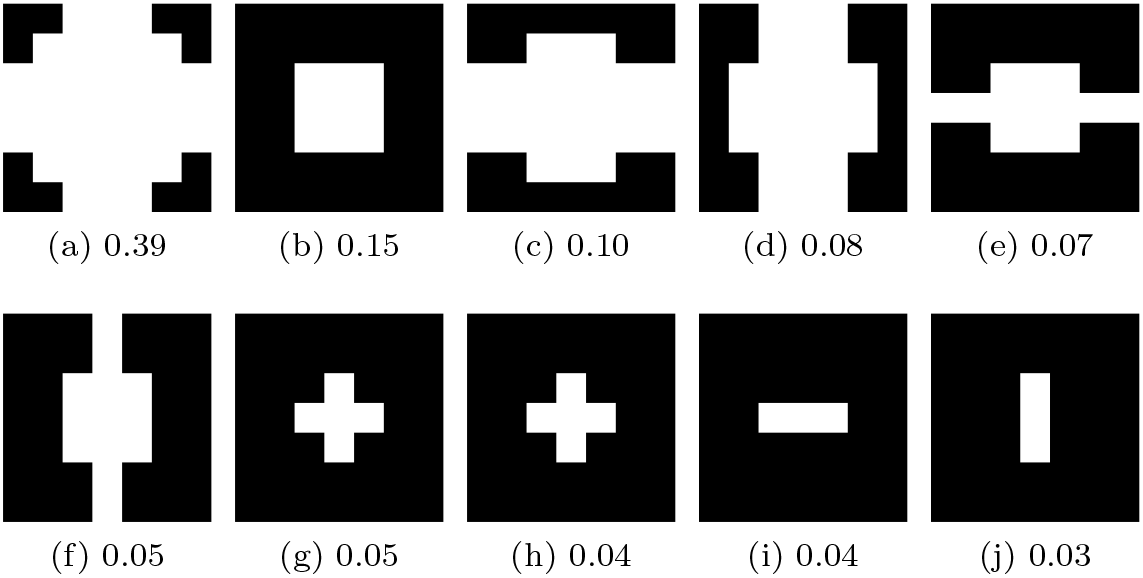
Phosphenes used in the model and their corresponding probability. These phosphenes are binary and 7 pixels wide. Images (h) and (i) represent the small oblong horizontal and vertical phosphenes, they are indistinguishable due to the low resolution.

**Figure 3:**
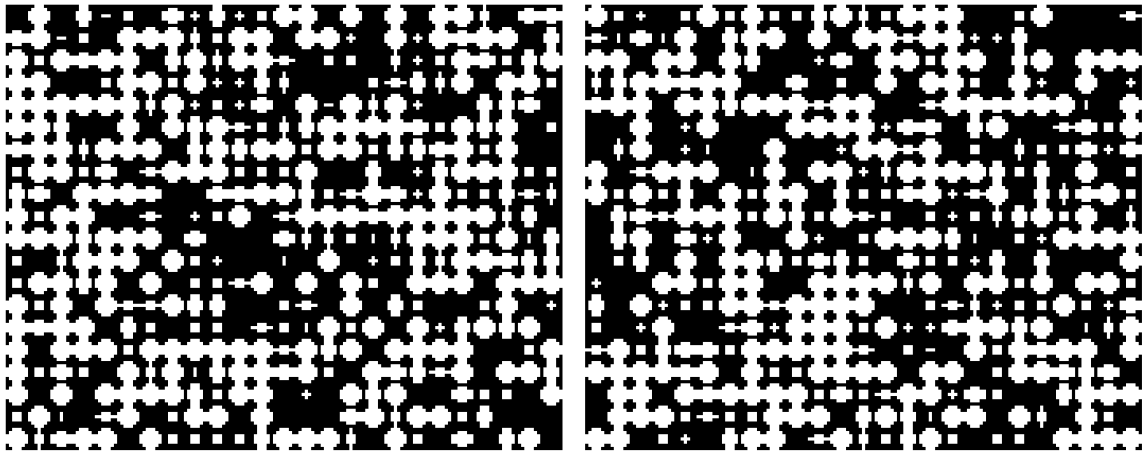
Typical phosphene maps / phosphene rendering for a full field stimulation. The phosphenes are shown with no overlap to be distinguishable from one another. In reality, the maps are used with an overlap of 4 pixels/ DMD mirrors. We also randomly not project phosphenes to simulate non perceived phosphenes.

## 3 Scanning behavior and head movements

In order to complete our phosphene model with a dynamic component, we measure the velocity of the head movements of an implanted patient. The statistics for the movement of the head was assessed based on an experiment conducted with a patient enrolled in clinical trials at the XV-XX Ophthalmological hospital in Paris.

### 3.1 OptiTrack system

The OptiTrack system^3^ is a motion capture system that outputs reliable values for the 3D pose of rigid bodies, provided that they are equipped with a number of infrared markers. Fixing markers on the object and the camera allows us to obtain their poses in the 3D space, from which we retrieve the pose of the object relative to the camera.

The cameras acquire the position of the infrared markers present in the scene at a frequency of 120 Hz. Prior to the acquisition, a number of *Rigid bodies* can be defined by selecting a subset of markers in the fields of view of the cameras. These markers are then tracked from one frame to the next, and the software directly outputs the position and orientation of the rigid bodies.

### 3.2 Setup

The patient was seated in front of a screen, in a dark room. The screen measures approximately 40 cm × 40 cm and was located at a distance of 35cm from the head of the patient. The task consists in finding white squares (7 cm × 7 cm) displayed on the screen over a dark background. The white square is displayed for as long as the patient needs it. The task ends when the subject touches the screen to indicate its response. On the set of approx. 20 trials that we recorded, it took between thirty seconds and two minutes to complete the trial. We did not record whether the responses were correct or not.

We placed three OptiTrack cameras behind the patient, at approximately two meters from the floor. The cameras were behind the patient and had the screen and the head of the patient in their field of view. Markers were set (i) on the head of the patient and (ii) on the top of the screen. Two rigid bodies were created in Motive^4^ (the software provided with OptiTrack) so that we could export directly data about these two rigid bodies. Three markers were set on the top of the screen and composed the screen rigid body. Eight markers were placed on the head, but only six of them were captured to define the head rigid body. The two remaining markers were recorded as well, but not used thereafter because the first six markers provided enough data to specify the motion of the head in all trials. Figure.4 shows the relative positions of the subject, the screen and the OptiTrack sensors.

A different take was created for each trial, providing 21 takes. Each take starts when the bright square appears on the screen and finishes after the subject has given its answer. Takes do not necessarily terminate exactly when the subject gives the answer. Hence the following analysis is realized during the time the subject is engaged in the scanning task.

**Figure 4:**
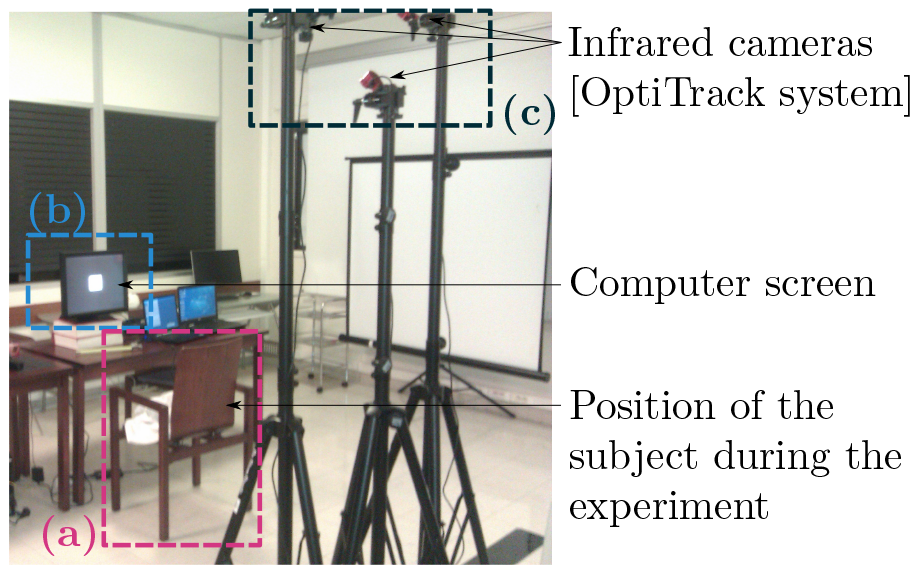
Experimental setup for the acquisition of head motion data. The subject performs a task while seated — location shown in (a) — in front of a computer screen (b). Three OptiTrack infrared sensors (c) are located behind the subject, at a height that puts both the subject’s head and the screen in their field of view.

### 3.3 Analysis

In the context of this study, the features of interest are the temporal dynamics of the head. In fact, as the camera is fixed on the goggles, the angular velocity of the head dictates the magnitude of the optical flow caused by the background and fixed objects on the array of photosensors of the camera.

Our work is based on the data exported from Motive. The software directly provides rigid bodies data. The orientation data is represented as quaternion. For each frame *n*, a quaternion 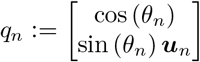 is provided. It represents the rotation with respect to a fixed initial position. *u*_*n*_ ∊ ℝ^3^ is a unitary vector representing the axis of the rotation, *θ*_*n*_ is the angle of the rotation.

The instantaneous rotation between two successive frames *n* and *n* + 1 is obtained by dividing *q*_*n*+1_ by *q*_*n*_. This operation provides a quaternion 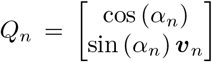. Denoting Δ*t* the time step associated with the sampling frequency of the Opti-Track sensors, the instantaneous angular speed is provided by *α*_*n*_/Δ*t*. Head movements were distributed along a mean angular velocity of 7.8 ± 5.4°/*s* during the task.

## 4 High temporal resolution stimulation platform

### 4.1 Participants

Ten normally sighted individuals, aged between 22 and 67 years (37.47 ± 15.9) participated in this study. Participants sight was controlled prior to the first experimental session to ensure that all participants acuity was superior to 8/10 (e.g <=0.1 logMAR), they also had to have a Minimental Status Examination Score superior to 27. This study was approved by the French ethics committee (CPP Ile de France IV). Before inclusion and after being informed of the experimental protocol each participant signed a consent form.

### 4.2 Experimental platform

A stimulation platform capable of rapidly switching between different temporal resolutions in a precise-control manner is required to perform a study on the effects of different frame rates (see Figure 5a). We used a Texas Instrument LightCrafter projector controlling a DLP3000 Digital Micromirror Device (DMD)^5^. The DMD comprises an array of mirrors reflecting light from a LED source, as described in figure 5b. Mirrors in the device can rapidly switch between two discrete angular positions (−12° and 12°) to enable or disable projection of light [23]. Mirror switching occurs at a very fast rate, so as to project different intensity levels using binary pulse-width modulation.

**Figure 5:**
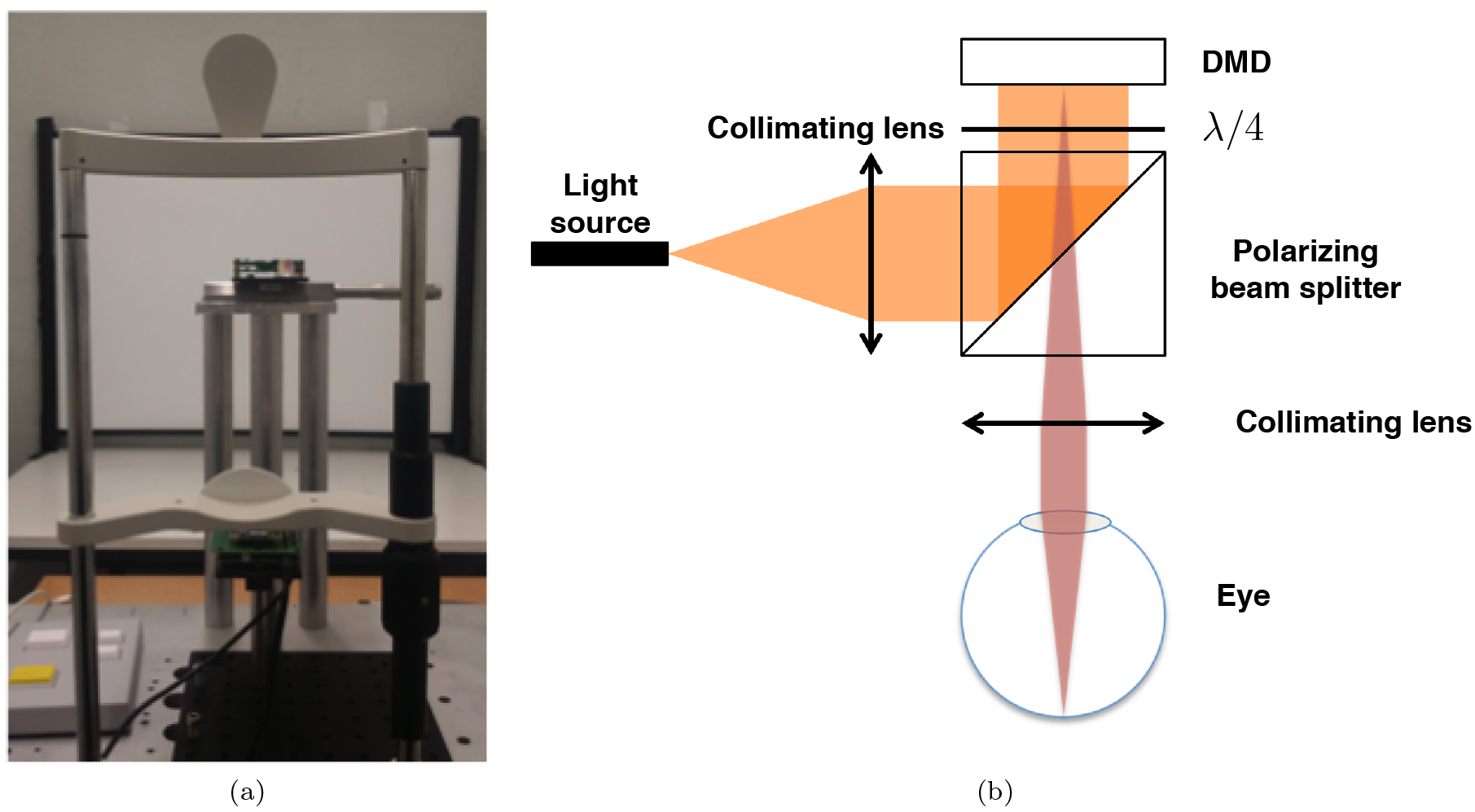
a) Picture of the experimental platform composed of the DMD and the chinrest. b) Description of the optical system used to project light directly into subjects’ eyes

In this work, the characteristics of DMD devices have been exploited to investigate the temporal properties of the visual system. The DMD was controlled with a real-time embedded Linux system based on an NVIDIA TK1 SoC, with bespoke software enabling the control of any mirror independently (e.g. no need to send the whole image) with a millisecond precision. In other words, binary stimulation up to 1 KHz frame rate can be projected.

The DLP3000 system can provide binary stimulation up to 1440 Hz at 100 % duty cycle. The platform allows stimulation display using different frame rates in a very rapid, bandwidth efficient way. The duration of the on-position for each mirror is configurable independently. This permits to keep it in a reflective state for a longer time (up to 694 ms) without resending any command to the device, significantly reducing the bandwidth required to operate the system.

Stimuli could be projected up to 1440 Hz in order to evaluate the optimal temporal resolution, nevertheless the perceived frame rate depended on the stimulus dynamic. For instance, even with an update of the visual information every 694 ms if the stimulus moved slowly with subpixel displacement every update the perceived frame rate would be lower than 1440 Hz as the visual information will change slower.

The platform was optically designed in order to project directly into the participant’s eye which provided an immersive stimulation. The optical design is described in figure 5b. Light power was limited at 5 *mW*/*mm*^2^ and controlled during the experiment to ensure security of the participants.

## 5 Experimental validation

### 5.1 Task I: Counting bars

In this first experiment, we measure the ability of observers to count the number of bars with different temporal resolutions in order to assess an acuity limit. The phosphene model obtained from 2.2 was used to provide a stimulation as close as possible to what implanted patients can see. For each trial a phosphene map was randomly generated with respect to the probability of appearance of each phosphene. Moreover, the dynamic of the stimuli was based on the velocity computed with the head movement data described in 3.

#### 5.1.1 Stimuli

The stimuli consisted of parallel vertical lines whose count was varied from 1 to 5, with 3 different spacings (from 0.44 deg to 0.86 deg) between them. Lines measured 5.9 deg in height and 0.11 deg in width and were flashed for 420 ms. Example stimuli for different distances and number of lines are presented in Figure 6(b) with their transformation according to our phosphene model. The experimental paradigm is outlined in Figure 6(a). The influence of stimulation duration was explore using three different temporal resolutions. Two 30 Hz stimulation with either a 100% or a 17% duty cycle and a high frame rate (HFR) condition where the projector was updated at a 1 KHz (which lead to a 200 Hz stimulation regarding to the stimulus speed). In the rest of the paper, the temporal condition at 30 Hz with 17% will be refereed as the flashing condition, and 30 Hz at 100% as “conventional” projection.

**Figure 6:**
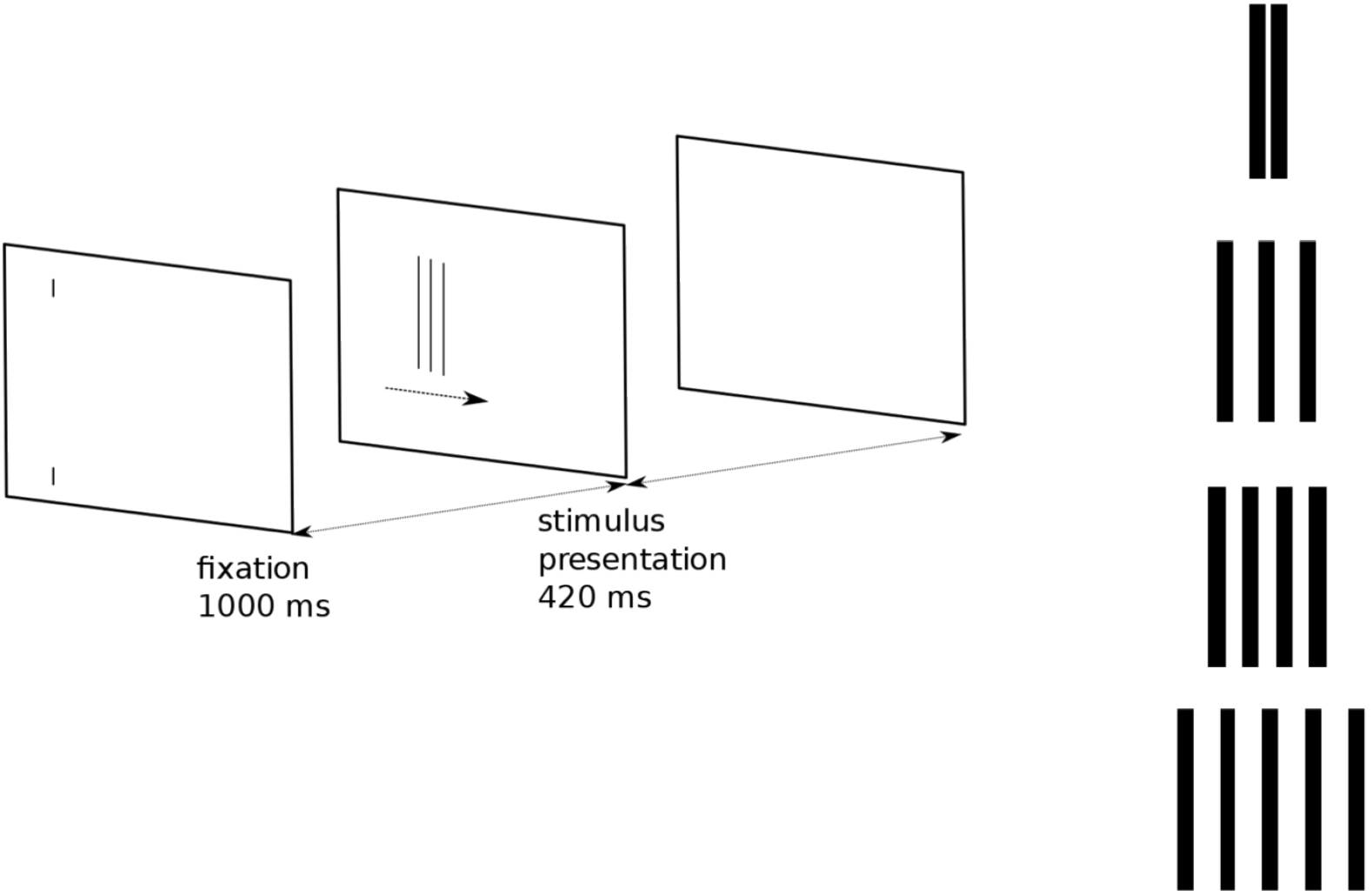
(a) Experimental paradigm: after 1*s* of fixation, a set of lines started moving horizontally across the screen for 420 ms. Participants were asked to report the number of line observed, and they were instructed to pursuit the stimulation. (b) Stimuli consisted in 2 to 5 vertical lines with different spacings.

Participants were instructed to first look at a the fixation point, once it disappeared they were ask to look at the stimulus and pursue it while it moved at a speed of 10 deg/s. At the end of each trial participants had to report how many bars appeared (between 1 and 5). Experimental conditions were then the following: 2 stimulus speeds *s*, 3 temporal resolutions *t*, 3 distances between bars *d*, 5 bar patterns *b* (with no distance condition for 1 bar) with 6 repetitions. This resulted in a total of 468 trials per subject.

#### 5.1.2 Results

Overall performance was better when stimuli were static (static: 79.6±1.46%, dynamic: 68.5±1.79%, F(1,700) = 23.17, *p* < 0.001). Nevertheless, participants had a lower success rate when comparing pulsed stimulation to HFR (pulse: 88.24±2.63%, HFR: 100±2.42%, Mann-Whitney U test *p* < 0.05). This result is mainly explained by the higher performances of participants with the HFR compare to the pulsed projection when the distance between bars was the smallest (HFR: 81.6±5.11%, pulse: 53.8±5.14%, Mann-Whitney U test *p* < 0.05).

Higher differences in performances were observed with moving stimulation between HFR (92.9±2.86%) condition compare to both conventional (76.4±3.04%, Mann-Whitney U test *p* < 0.05) and pulsed projection (73.7±3.3%, Mann-Whitney U test *p* < 0.01).

Discrimination is lower when considering trials at the smallest distance (0.44 deg) between bars. Performances fall dramatically below chance level for the pulsed condition (16.2%± 5.27), results at the HFR (56.32%±5.75) also decrease but are still significantly high (Mann-Whitney U test *p* < 0.01). In the same distance conditions, results with conventional projection (34.8%±5.46) are also lower than in the HFR condition, nevertheless statistical analysis show no significant different. For the middle distance condition, performances are significantly higher between HFR (100±3.91%) and both conventional (77.8±4.17%, Mann-Whitney U test *p* < 0.01) and pulsed projection (82.6±4.24% Mann-Whitney U test *p* < 0.01). These results are summarized figure 7.

**Figure 7:**
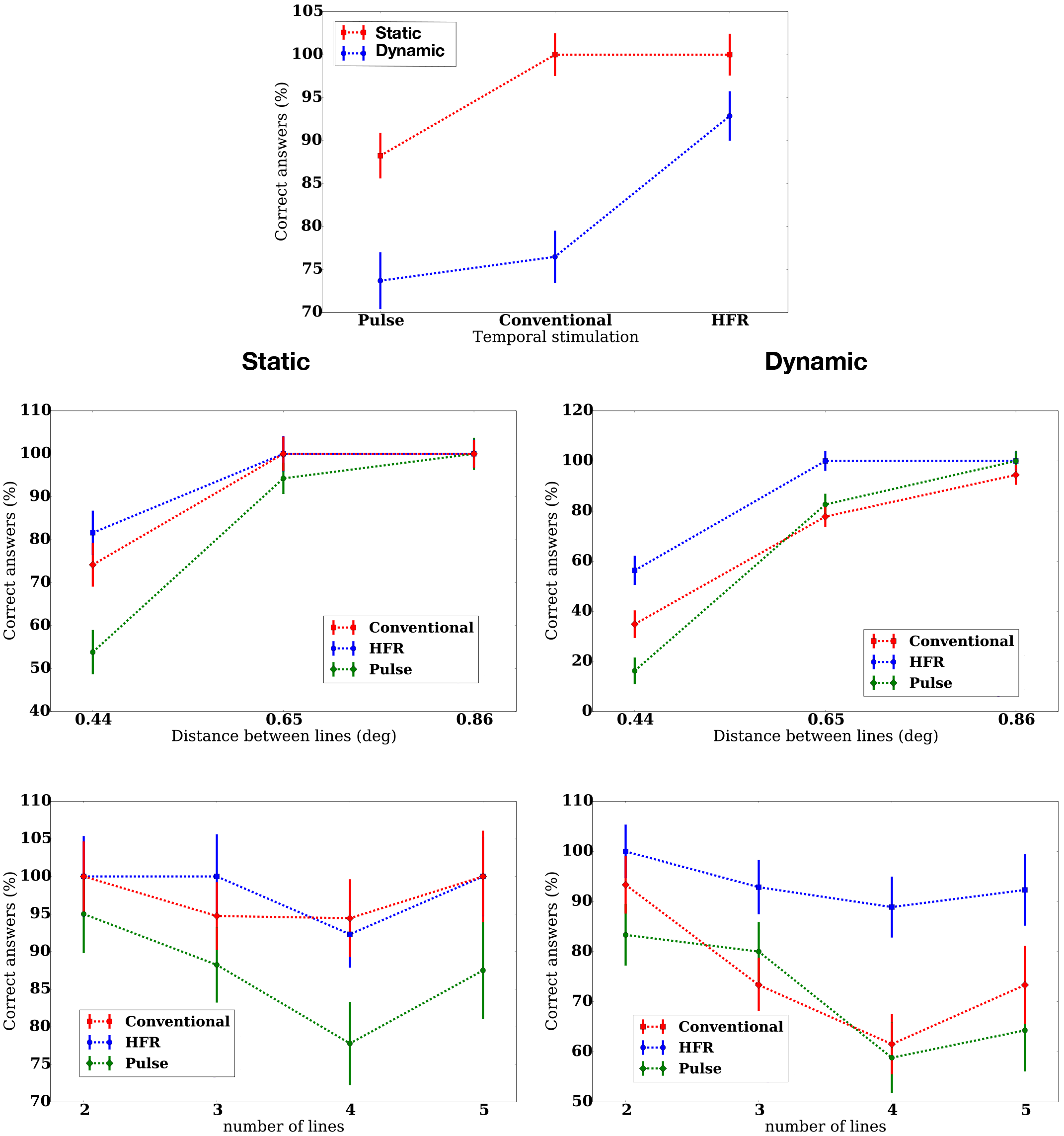
Overall results for the counting line experiment. Top figure represent the percentage of correct answer in function of stimulus dynamic and frame rate. Left and right panels, respectively static and dynamic conditions, present results with the different frame rates for different spacing and number of lines.

### 5.2 Task II: Numbers recognition

In the second experiment, the ability to discriminate more complex patterns was evaluated. This task provides information on the performance of patients with prosthetic vision when reading digits or characters.

#### 5.2.1 Stimuli

The stimuli were composed of three digits, two of the digits were ‘8’ and the third one which was the one to discriminate could be any other number (it could not be ‘8’). The position of the digit to recognize was randomly chosen across trial (see Figure.8).

**Figure 8:**
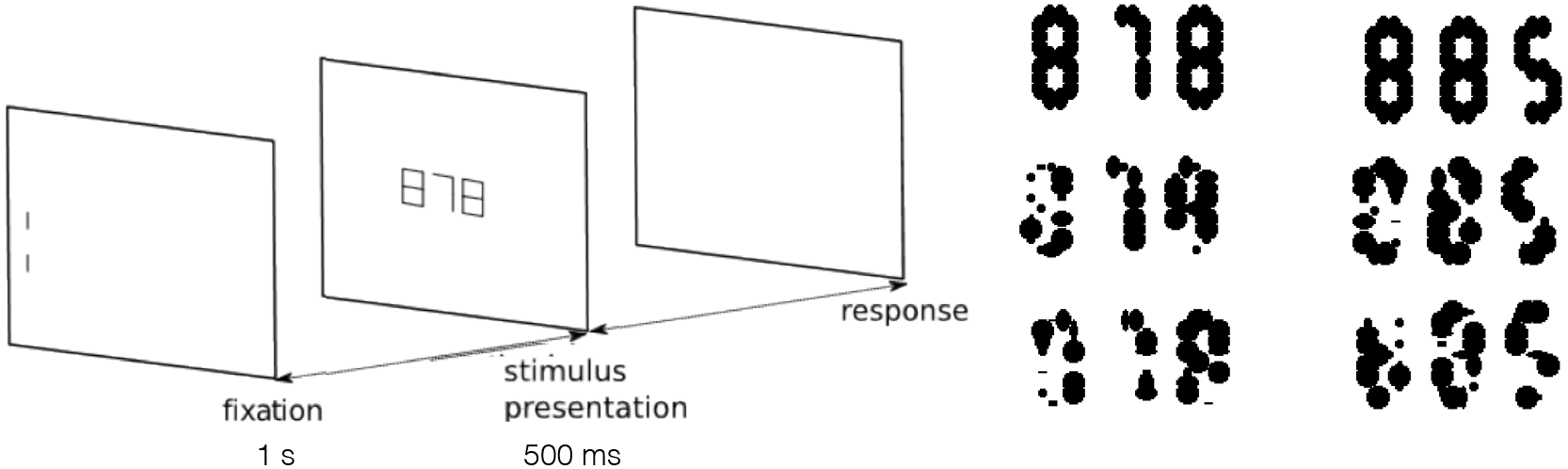
Digit identification experiment: (a) after an initial fixation cue of 1 s, the digits appear and move for 500 ms. Participants are instructed to track the stimulus and to report which digit was presented after the trial ends. (b) Stimuli with perfect phosphene map (top) and examples of the same digits using our phosphene model. Two spatial separations between digits were used: 0.10 deg and 0.32 deg.

As in the bar-counting experiment, the stimulus appeared after a fixation for 500 ms and was either static or moved at 10 deg/*s*. The impact of distance between digits on performance was also evaluated with 2 different separation distances (0.10 deg and 0.32 deg). Temporal resolution varied according to the first experiment with the same frame rate and duty cycle (HFR, 30 Hz and flashing). At the end of each trial participants reported to the experimenter which number they identified.

#### 5.2.2 Results

Comparison of performance between static and dynamic stimuli gave the opposite results of those obtained in the first task, as describe in Fig 9A. Overall, participants’ performance increased of about 20% when the stimulus was moving (static: 55.7±2.19%, moving: 74.8±2.50%, F(1,286) = 32.53, *p* < .001.)

**Figure 9:**
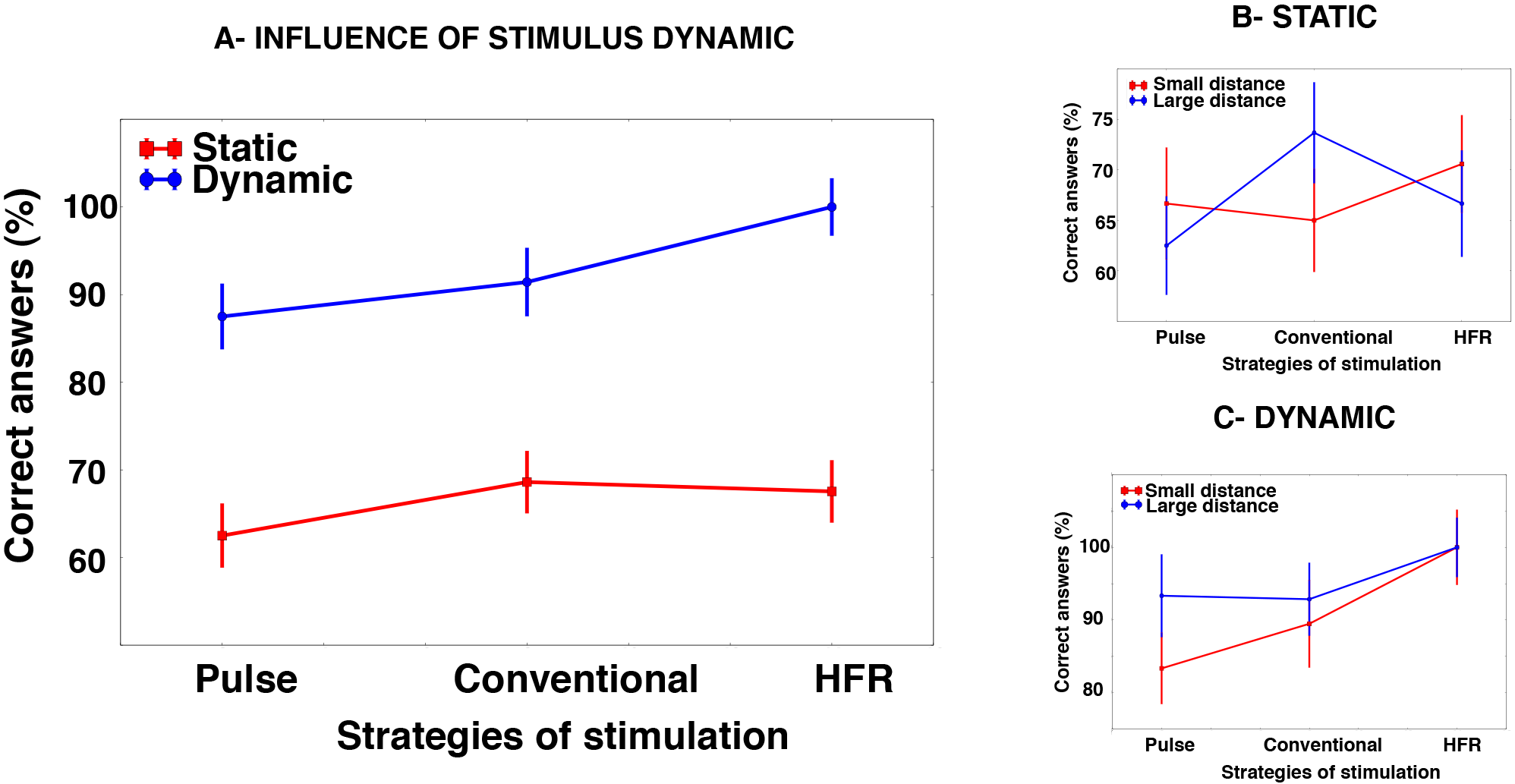
Performances during numbers recognition task: A) Discrimination for static (red) and dynamic (blue) stimuli is represented in function of the different stimulation rate (pulse, conventional, HFR), performance was also evaluated for the static (B) and dynamic (C) stimulation with a small (red) and large (blue) distance between digits.

Considering only data with static stimulus (Figure.9B), analysis of performance show no significant impact of frame rate (HFR: 65.7±3.64%, conventional: 62.5±3.93%, pulse: 62.0±3.88%), nor of distance between numbers (small: 63.8±3.07%, large: 62.5±3.15%). Improvement of discrimination performances with dynamic stimulus (Figure.9A) was significantly higher with the HFR condition by comparison to both conventional (Mann-Whitney U test, z-score = −3.45, *p* < .001) and pulsed conditions (Mann-Whitney U test, z-score = −3.78, *p* < .001).

As described in Figure.9C, the HFR condition was not affected by the size of the digit separation, even considering the smallest distance. Performances with the HFR projection were still high and significantly better than with the conventional (Mann-Whitney U test, z-score = −2.08, *p* < .05) or pulsed projection (Mann-Whitney U test, z-score = −2.41, *p* < .05).

## 6 Discussion and Conclusions

The quality of vision restoration provided by retinal implants were sought to be determined by the number of electrodes. This study shows that spatial resolution is not the only factor enhancing visual stimulation as increasing the temporal resolution of the stimulation improves visual perception in implanted patients, particularly when dealing with dynamic environments. Considering the dynamic dimension is essential for mobility as it is the most important component for the quality of life of visual impaired patients. We chose velocities range recorded from Retinitis Pigmentosa implanted patients performing head scanning movements. These head movements are developed by patients to explore environments and replace saccadic motion of eyes. Pixium’s implant imposes to fixate a fixed target stimulation forbidding any eye saccadic exploration movement that is then replaced by head movements. We also relied on phosphene models based on the description made by implanted patients of their perception to multiple single-electrode electric stimulation. Phosphene maps relied on the same shapes, sizes and distributions of phosphenes described by implanted patients. This allowed us to emulate the percepts induced by a retinal implant, and test our hypothesis about temporal resolution in healthy subjects. Results show an improvement of visual perception with the increase of temporal resolution which is consistent with previous studies of subjects with normal vision [19, 24, 25, 26]. We also found an impact of stimulus speed, with lower performance on discriminating dynamically moving lines, as compared to static lines, during the counting task. Conversely, during the digit recognition task participants achieved higher discrimination levels when stimulation digits were moving. This observation can be explained by the complexity of shapes in both tasks. High frame rates increased available information by spatially subsampling the stimulation by comparison to both conventional and pulsed projection. During the dynamic condition at HFR, the stimulus was refreshed 3 times more than with the conventional projection. While the line counting task provides an evaluation of the expected acuity with such a device, the digit discrimination task includes more complex stimuli, which are more likely to be observed in a natural environment and clearly shows an increase in pattern discrimination.

Increasing the frame rate to match the retina known temporal precision is providing somehow an expected result [16]. There as converging evidence from neuroscience suggests that neurons in early stages of sensory processing in primary cortical areas, including both vision and other modalities, use the millisecond precise time of neural responses to carry information. We believe that the widespread use of low temporal resolution is unrelated to the biology of the visual system [16][27], but is rooted in the visual depictions found in art and photography extending back to the origins of painting and is therefore optimal for reproducing high quality movies, rather than for extracting information, as needed in artificial vision.

In the brain, there has been evidence for a high precision of neural coding in different sensory structures ([27][28][29][30][31]). In the recognition task, objects were animated with a global jittering motion that bears some resemblance to fixational eye movements. The role of such eye movements in maintaining a high acuity in object recognition has been demonstrated in a recent study [32]. This suggests that the brain must use the precise timing frequencies of the retinal flow to optimize pattern recognition. However, the role of millisecond precise timing in encoding invariant patterns or objects to recognize objects by higher cortical areas has not been established yet [33]. There is however increasing evidence that, unlike what was initially believed [34], the precision of spike timing is not progressively lost as sensory information travels from the peripheral representation to the higher level areas closer to decisions and motor acts [35]. This therefore suggests that primary feature layers based on spike timing like the ones proposed in our model may be relevant for processing.

More interestingly there exists a technology allowing to acquire such fast visual information [36]. Inspired by the biology of the eye and brain, the neuromorphic field developed sensors containing arrays of independently operating pixel sensors that are now available off the shelves (and currently in use by Pixium) [36]. In these sensors, also referred to as silicon retinas, each pixel is attached to a level-crossing detector and a separate exposure-measurement circuit. For each individual pixel, the electronics detect when the amplitude of that pixel’s signal reaches a previously established threshold above or below the last-recorded signal level, at which point the new level is then recorded. In this way every pixel optimizes its own sampling depending on the changes in the light it takes in. Like biological retinas, the artificial sensors used in this work do not encode static images, but instead transmit incoming information instantaneously. Both are entirely asynchronous, biological retinas exhibit precision to the order of one millisecond, while event-based sensors are precise to the order of one microsecond. This precision can be used to derive precise timed models of the retina [37].

## Aknowledgment

Authors are greatfull to V.Bismuth and R. Hornig from Pixium Vision for their feedback and collaboration during this work.

## Footnotes

1 http://www.pixium-vision.com/file_bdd/dynamic_content/file_pdf_pdf_en/1456851776_PixiumVision-IRISIIActivation-ENGFinal.pdf

2 http://bionicvision.org.au/eye

3 http://www.optitrack.com/

4 http://www.optitrack.com/products/motive/

5 www.ti.com/tool/dlplightcrafter, last accessed 17/02/2015

